# Delta- and theta-band cortical tracking and phase-amplitude coupling to sung speech by infants

**DOI:** 10.1101/2020.10.12.329326

**Authors:** Adam Attaheri, Áine Ní Choisdealbha, Giovanni M. Di Liberto, Sinead Rocha, Perrine Brusini, Natasha Mead, Helen Olawole-Scott, Panagiotis Boutris, Samuel Gibbon, Isabel Williams, Christina Grey, Sheila Flanagan, Usha Goswami

**Affiliations:** University of Cambridge; Ecole Normale Superieure

**Keywords:** EEG, Language, Neural Oscillations, TRF, Infant

## Abstract

The amplitude envelope of speech carries crucial low-frequency acoustic information that assists linguistic decoding at multiple time scales. Neurophysiological signals are known to track the amplitude envelope of adult-directed speech (ADS), particularly in the theta-band. Acoustic analysis of infant-directed speech (IDS) has revealed significantly greater modulation energy than ADS in an amplitude-modulation (AM) band centered on ∼2 Hz. Accordingly, cortical tracking of IDS by delta-band neural signals may be key to language acquisition. Speech also contains acoustic information within its higher-frequency bands (beta, gamma). Adult EEG and MEG studies reveal an oscillatory hierarchy, whereby low-frequency (delta, theta) neural phase dynamics temporally organize the amplitude of high-frequency signals (phase amplitude coupling, PAC). Whilst consensus is growing around the role of PAC in the matured adult brain, its role in the *development* of speech processing is unexplored.

Here, we examined the presence and maturation of low-frequency (<12 Hz) cortical speech tracking in infants by recording EEG longitudinally from 60 participants when aged 4-, 7- and 11-months as they listened to nursery rhymes. After establishing stimulus-related neural signals in delta and theta, cortical tracking at each age was assessed in the delta, theta and alpha [control] bands using a multivariate temporal response function (mTRF) method. Delta-beta, delta-gamma, theta-beta and theta-gamma phase-amplitude coupling (PAC) was also assessed. Significant delta and theta but not alpha tracking was found. Significant PAC was present at all ages, with both delta and theta -driven coupling observed.

**Graphical abstract:** 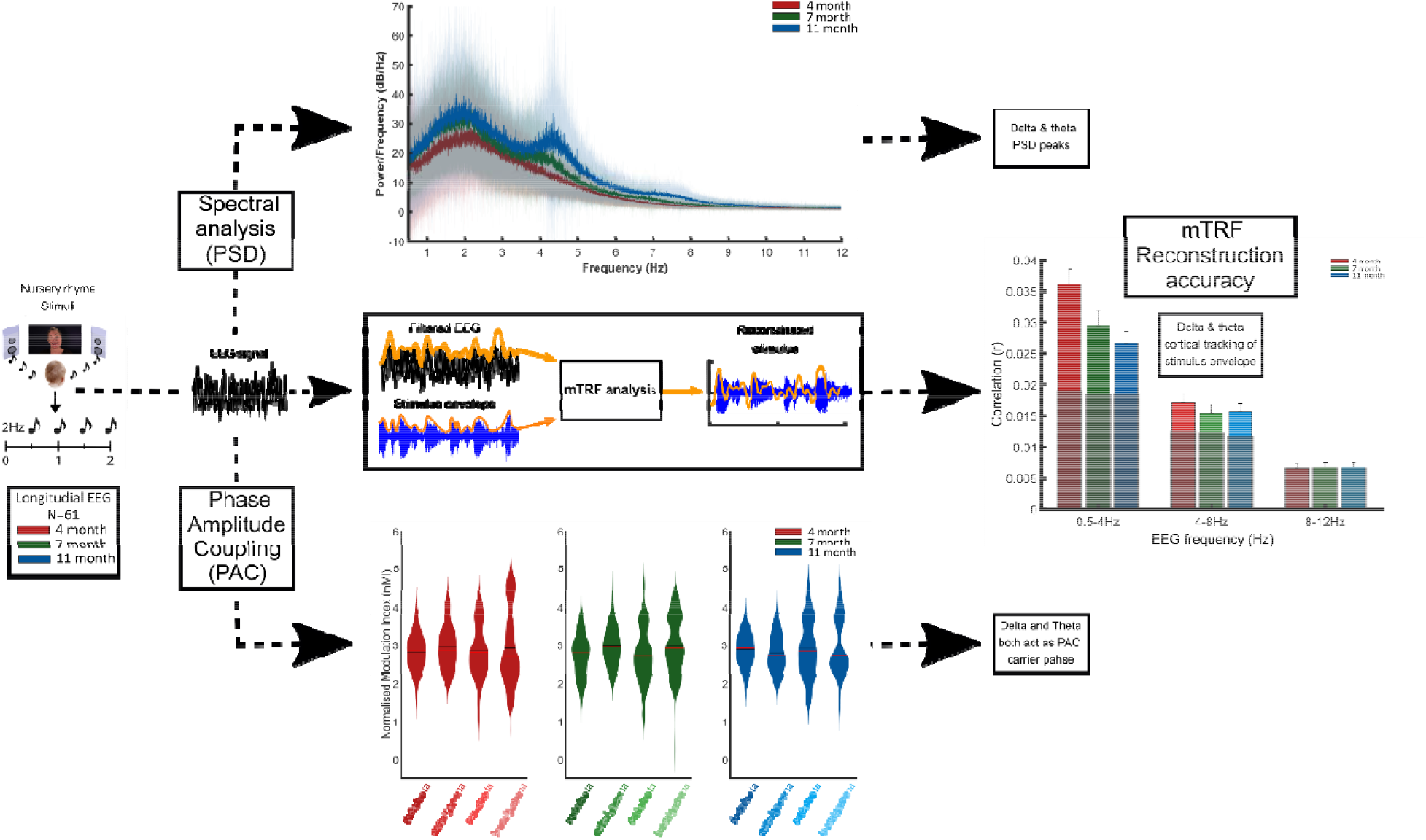

**Highlights:**

- Longitudinal EEG study in which 4, 7- & 11-month infants listened to nursery rhymes
- We demonstrate cortical speech tracking via delta & theta neural signals (mTRF)
- Periodogram (PSD) analysis revealed stimulus related delta & theta PSD peaks
- Delta and theta driven phase amplitude coupling (PAC) was found at all ages
- Gamma frequency amplitudes displayed stronger PAC to low frequency phases than beta

## 1 Introduction

Neural signals synchronise to the auditory envelope during speech listening. This cortical tracking phenomenon has an important functional role for speech comprehension in adults (1, 2). The speech envelope is the low-frequency amplitude modulation that carries crucial acoustic information required for perceptual encoding and linguistic decoding at multiple temporal scales. Theorists in speech perception have argued for special roles for syllable-level acoustic rhythmic information (slower temporal modulations centred on ∼5 Hz) and phoneme-level acoustic information (rapid temporal modulations centred on ∼ 40 Hz) (3). It is proposed that intrinsic neural signals in cortex at matching frequencies (theta, 4-8 Hz; low gamma, 30-50 Hz) provide one mechanism for encoding this acoustic information, parsing speech signals into temporal chunks or windows for subsequent analysis and comprehension (multi-time resolution (2)). Delta-band tracking in the adult brain is also found, thought to reflect discourse-level tracking and perceptual grouping (4, 5). Low-frequency (delta, theta) phase and high-frequency (gamma) amplitudes also track acoustic rhythms in the speech signal by operating together as an integrated representational mechanism via phase-amplitude coupling, PAC (6–8), where the phase dynamics of low-frequency neural signals temporally organize the amplitude of high-frequency signals (9–11). Adult studies have demonstrated the functional importance of PAC for speech encoding, with delta and theta phase coupling with gamma amplitude during stimulus-specific speech sampling (6). The role of cortical tracking for speech intelligibility has also been demonstrated by EEG (12), MEG (13–15) as well as intra-cranial measurements in adult patients using electrocorticography (ECoG) (16–18). Studies measuring adult cortical tracking of unintelligible speech report reduced tracking compared to intelligible speech (5, 6, 12–15, 19– 21), however the data are mixed, with the role of cortical tracking to the different elements of speech, and their role in speech intelligibility widely debated (12, 22–26). To date almost all available studies of PAC and cortical tracking at different frequencies rely on adult neural imaging (21), indicating a reasonable degree of consensus concerning neural mechanisms, with a core role established for theta tracking (5, 12, 14, 21).

While theta tracking may be a core feature of adult speech encoding, it is important to remember that the adult participants in neural studies have years of listening experience. Accordingly, the theoretical weight given to theta and gamma oscillations and linguistic units (2) may reflect top-down processing and the end state of development. Studies of children suggest that the weighting of different speech encoding mechanisms may change with development (27). For example, data from children with developmental dyslexia show core cognitive impairments in processing phonological (speech-sound) information, and associated atypical neural synchronisation to, and tracking of, speech input in the *delta band*, not in the theta band (4, 12, 28–30). Regarding degraded speech, typically-developing children show delta-band tracking of speech-in-noise, but not efficient theta-band tracking (31).

In adult studies, delta-band tracking is frequently related to auditory attentional mechanisms linked to the processing of sequential attributes of speech sounds, for example grouping (21, 32), and to discourse-level parsing related to phrasing (14, 33). While the pre-verbal infant brain may also utilize grouping mechanisms, infants have not yet learned discourse-level speech information (34). Cross-language linguistic data show that strong syllables are produced approximately twice per second (2 Hz) in all languages, both stress-timed and syllable-timed (35). Strong (stressed) syllables may provide a set of acoustic landmarks at a delta-band rate that facilitate grouping, enabling the infant brain to begin to represent the acoustic structure of human speech. Even newborn infants can discriminate languages from different rhythm classes (stress timed [e.g. English], syllable timed [e.g. French] or mora timed [Japanese]) (36). Rhythm perception has thus been described in the cognitive literature as a language-universal precursor to language acquisition (37). One way to study early cortical tracking of speech is to use sung or chanted speech, as exemplified by the nursery rhymes of the English cradle (38). Further, the role of PAC in the *development* of speech processing by children and infants is not yet explored.

There are three previous infant language-based cortical tracking studies of speech, two with 7 month-old infants (39, 40) and one with neonates and 6-month-olds (41). These prior studies were not longitudinal and adopted either a forward linear modelling approach (detailed methods; 37), or a cross-correlation analysis, to estimate how well the amplitude envelope of IDS was represented in the cortical activity. All studies reported significant cortical tracking, however none addressed the potential developmental roles of delta-versus theta-band tracking regarding language processing, nor examined possible developmental changes for delta versus theta tracking. To date, there are two PAC studies in infants. Both have focused on clinical applications, specifically the role of PAC in functional connectivity (42, 43). For example, one study measuring PAC in sleeping newborns during active versus quiet sleep reported hierarchical nesting of oscillations in both sleep states (42). Accordingly, current data suggests that PAC mechanisms in the infant brain may be online at birth. However, no study to date has looked at the functional relevance of infant PAC when processing sensory stimuli, leaving PAC during language processing yet to be explored.

Here we assess infant cortical tracking of the temporal modulation information in IDS present in both delta- and theta-rate AM bands. We analysed cortical tracking of nursery rhymes, originally recorded being sung or chanted to an alert infant. We used sung speech as theoretically it provides an optimally-structured stimulus for the infant brain to employ cortical tracking of speech, since all the amplitude modulations at different frequencies are temporally aligned. Although the center frequencies of canonical EEG bands may shift during infancy (44, 45), the delta band is relatively stable, and there is no consensus on the exact boundaries of different EEG bands at specific time points in development (35). There are also no infant data regarding center frequencies from studies utilizing speech stimuli. Accordingly, the EEG delta and theta bands were defined following the prior speech processing literature (delta, 0.5-4 Hz; theta, 4-8 Hz). These bands have also been identified by modelling the temporal modulation architecture of IDS (46). Leong et al. theorised that the amplitude envelope of spoken IDS contains a nested oscillatory hierarchy of amplitude modulations (AMs) in three temporal bands, corresponding to the timing of delta, theta and beta/low gamma cortical neural signals. Furthermore, they demonstrated that the low-frequency AM bands in IDS showed different degrees of coupling with each other compared to adult-directed speech (ADS). IDS both had significantly more modulation energy in the delta band than ADS (which had significantly more modulation energy in the theta band), and significantly greater phase synchronisation between the delta-rate and theta-rate bands of AM than ADS. Based on this computational modelling (46), we therefore hypothesised that delta-band neural signals would track the speech envelope most strongly early in infant development. We also tentatively hypothesised that theta tracking may come online more slowly.

64-channel EEG was recorded from a sample of 60 infants as they watched and listened passively to a video of a range of nursery rhymes with different beat structures being sung or rhythmically chanted by a female “talking head”. The video lasted approximately 20 minutes (see Figure 2a), and was followed by a resting “silent state” recording. Prominent modulation spectrum peaks were identifiable in the nursery rhyme stimuli at rates corresponding to adult delta and theta neural signals (Supplementary Figures S2, S3 and S4). The neural data presented here were recorded from the first half of infants recruited to an ongoing longitudinal study of 113 infants (122 infants recruited, 113 retained), who listened to nursery rhymes at the ages of 4-, 7- and 11-months. Data collection with the other infants is still ongoing. Our approach allows for subsequent confirmatory testing of the current results with the second half of the sample.

Multivariate temporal response functions (mTRFs) were employed to measure cortical tracking (47). mTRFs are encoding models that describe how the inputs and outputs of a system are linearly related. Forward models provide researchers with information regarding the precise spatial and temporal patterns reflecting the neural processing of a stimulus, as used previously with infants (30, 31). A backward ‘stimulus reconstruction’ version of the TRF linear modelling technique, not yet used with infants, was employed here. The backward mTRF uses information from all EEG channels simultaneously to reconstruct the speech envelope, thereby enabling increased sensitivity to signal differences between highly correlated response channels, thus offering several advantages (47) over forward modeling (39, 40) or cross-correlation approaches (41). The mTRF provides a single summary measure of speech-EEG coupling for data recorded over long periods of time by assessing the fidelity of the reconstruction (47). Here, we reconstructed the heard speech in the delta and theta bands, using stimulus reconstruction in the alpha band as a control condition. Although alpha band responses may be important in adult speech tracking (48–50), alpha-band AMs did not play a role in the prior computational modelling of IDS (46). Accordingly, we do not predict significant cortical tracking in the alpha band for the infant brain.

Finally, we hypothesised that low-frequency oscillatory phase, specifically within the delta band, would couple with high-frequency amplitudes in the infant brain during speech processing. Given our hypothesis that delta-band neural signals would be particularly prominent early in infant development, we predicted that delta-driven PAC should emerge prior to theta-driven PAC. If individual differences in sensory processing drive cognitive development rather than individual differences in brain mechanisms (51), the basic neural mechanisms for speech encoding should be present at birth.

## 2 Materials and Methods

### 2.1 Infant participants and ethics

Data from the first half (N=60) of a larger longitudinal cohort (122 infants recruited, 113 retained) were used in all analyses. From this initial cohort, four participants withdrew without providing any data and one withdrew after the first session so was also excluded. The remaining sample of 55 infants provided data at ∼4-months (115.6 □ ± 5.3 days), ∼7-months (212.2 □ ± □ 7.2 days) and ∼11-months (333.1 ± 5.6 days) [mean ± standard deviation (*SD*)]. Due to missed appointments (7-months N=2), technical issues with the data file (4-months, N=1) the final number of data points at each month was, ∼4-months (N=54), ∼7-months (N=53), ∼11-months (N=55).

Infants were recruited from a medium sized city in the United Kingdom and surrounding areas via multiple means including flyers in hospitals, schools and antenatal classes, research presentations at maternity classes and online advertising. All infants were born full term (37-42 gestational weeks) and had no diagnosed developmental disorder. The study was reviewed by the Psychology Research Ethics Committee of the University of Cambridge. Parents gave written informed consent after a detailed explanation of the study and families were repeatedly reminded that they could withdraw from the study at any point during the repeated appointments.

### 2.2 Stimuli

A selection of 18 typical English language nursery rhymes were chosen as the stimuli. Audio-visual stimuli of a singing head were recorded using a Canon XA20 video camera at 1080p, 50fps and with audio at 4800 Hz. A native female speaker of British English used infant directed speech to melodically sing (for example “Mary Mary Quite Contrary”) or rhythmically chant (for nursery rhymes like “There was an old woman who lived in a shoe”) the nursery rhymes whilst listening to a 120 bpm metronome. Although the nursery rhymes had a range of beat rates and indeed some utilized a 1 Hz rate (Supplementary Figures S2 and S4), the metronome was used to keep the singer on time. The beat was not present on the stimulus videos, but it ensured that a consistent quasi-rhythmic production was maintained throughout the 18 nursery rhymes. To ensure natural vocalisations the nursery rhyme videos were recorded sung, or rhythmically chanted, live to an alert infant or sung again immediately afterwards if the infant fussed out.

### 2.3 EEG data collection

Infants were seated in a highchair, approximately one meter in front of their primary care giver, within a sound-proof acoustic chamber. EEG data was recorded at a sampling rate of 1000 Hz using a GES 300 amplifier connected to a correctly sized 64 channel infant electrode net (Geodesic Sensor Net, Electrical Geodesics Inc., Eugene, OR, USA). The infant was seated ∼650mm away from the presentation screen and sounds were presented at 60dB (checked by sound level meter) from speakers (Q acoustics 2020i driven by a Cambridge Audio Topaz AM5 Stereo amplifier) placed either side of the screen. Whilst the infant attended to the screen, the 18 nursery rhyme videos played sequentially, each repeated 3 times (54 videos, with a presentation time of 20’ 33’’ in total). If the infant lost attention to the screen an attention grabber video was played at the end of that nursery rhyme. If the infant became too fussy a short break was allowed before continuing, otherwise the session was ended. All infants included in analysis listened to at least 2 repetitions of each nursery rhyme (minimum of 36 nursery rhymes, lasting 13’ 42’’). This stimulus period was followed by 5 minutes of silent recording (silent state). To ensure compliance from the infant, and therefore a good EEG signal during the silent state, it was necessary for a researcher to sit alongside the infant during the silent state recording. To ensure consistency across participants, the researcher performed the same action of silently blowing bubbles and showing the same picture book to the infant.

### 2.4 EEG preprocessing

All analyses were conducted with custom-made scripts in Matlab 2017a (The MathWorks, Inc., Natick, MA) incorporating the EEGLab toolbox (52).

The four (channels 61, 62, 63 and 64) facial electrodes were excluded from all analyses because they do not exist on the infant-sized EGI Geodesic sensor nets. The EEG data, from the remaining 60 channels, was filtered (*pop_eegfiltnew* function of EEGLab toolbox) into a broadband signal (0.5-45 Hz for PSD and PAC, methods for acquiring PAC bands of interest further detailed in PAC methods) or the frequency band of interest (0.5-4 Hz, 4-8 Hz or 8-12 Hz for the mTRF analysis) using zero-phase bandpass Hamming windowed FIR filters (transition band widths of 2 Hz with cutoff frequencies at -6 dB, 0-5 Hz, 3-9 Hz and 7-13 Hz respectively). The EEG data was down sampled to 100 Hz to reduce the computational load. Next, the *clean_asr* EEGLab function (52) was used to clean noise artifacts from the data by identifying and removing bad principal components via a modified PCA procedure (see supplement for more details). Further bad channels were identified via probability and kurtosis and were interpolated (via spherical interpolation), if they were 3*SD* away from the mean (average number of interpolated channels; 4mo = 6.7, 7mo = 6.3, 11mo = 6.6), before all channels were re-referenced to a 60-channel average reference.

EEG responses were epoched into trials aligned to the start and ending at the completion of a phrase (e.g. “Mary had a little lamb”), producing EEG responses to 83 phrases (*M length* ± *SD*: 4.23sec ±0.88) which were repeated a maximum of 3 times in the experiment (249 epochs in total). This epoching procedure was decided as infant EEG typically contains short, irregular, movement artifacts and using shorter epochs increases the likelihood of generating a whole artifact free epoch whilst maintaining long enough epochs for optimising the model fit with the mTRF toolbox (47). The infants occasionally rested their head or touched the EEG cap in such a way that specific channels showed short bursts of noise whilst the rest of the channels remained clean. To account for this, and retain a large volume of the data, epoch by epoch channel rejection was conducted. Per epoch, probability and kurtosis were used to identify bad channels and were interpolated (via spherical interpolation) if they were 3*SD* away from the mean. Finally, bad epochs were rejected with the *pop_autorej* function (EEGLab), removing epochs with fluctuations above 1000uV and values outside a 3*SD* of the probability threshold.

### 2.5 Multivariate Temporal Response Function (mTRF)

TRFs are encoding models that can describe how an input and output of a system are related via linear convolution (47). Here, we applied TRFs in a backward direction to assess how strongly a stimulus property, in this case the stimulus envelope, is encoded in the neural response. The backwards, “stimulus reconstruction”, mTRF model has important advantages compared to previous forward linear encoding models in infants (39, 40). Specifically, it utilises data from all the EEG channels simultaneously in a multivariate manner, while forward TRF models are fit on individual channels separately.

After preprocessing, the EEG responses to each of the nursery rhymes were averaged (data across a maximum of three repeats, of the 83 nursery rhyme trials collected, were averaged), thereby creating 83 “averaged trials”. This averaging was conducted to improve the signal to noise ratio of the data for the mTRF analysis (4-month, *75*.*9* ± 13.3 epochs; 7-month, *79*.*4* ± 6.4 epochs; 11-month, *80*.*7* ± 5.1 epochs [mean ± standard deviation (*SD*)]). If less than half (42 averaged trials) remained after preprocessing the file was removed from further analysis (minimum numbers at 4-month, 47 epochs; 7-month, 47 epochs and 11-month 65 epochs). The mTRF analysis was conducted using the multivariate temporal response function (mTRF) toolbox (47) through Matlab 2017a (The MathWorks, Inc., Natick, MA). The backwards model can be expressed by the following formula in which the reconstructed stimulus envelope ŝ*(t)* is created by a linear decoder, g(τ, n), mapping the neural response, *r(t,n)*, back to the stimulus, *s(t)*. The TRF decoder was used to integrate the neural responses at multiple time lags, τ, between 0 and 250ms (τmin = 0ms, τmax = 250ms). These ‘stimulus-relevant’ time lags where selected in keeping with the previous literature (5, 28, 47)

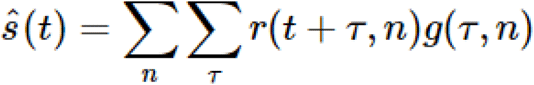

The quality of the envelope tracking within each EEG frequency band was assessed by a “leave-one-out” cross-validation per infant (47). First the average trial EEG epochs (maximum of 83) were normalised via function *nt_normcol* (Noisetools *http://audition.ens.fr/adc/NoiseTools/*). Normalisation decreased the range of values that were necessary for the regularisation parameter search in the mTRF toolbox, making the cross validation more efficient. Next, the normalised average trials were rotated M-1 times, each serving once as the “test set” with the remainder of the trials being the TRF “training set”. For each rotation, the resultant M-1 training models were averaged to create one average model from the training set. The average model was subsequently convolved with the test data to reconstruct the stimulus. Pearson’s correlation (r) was used to validate how well the reconstructed stimulus envelope correlated to the original. This process was repeated for the M-1 rotations (maximum of 83 rotations if the data set had the full 83 averaged trials remaining after preprocessing). To avoid overfitting the model to a specific trial, an individual’s average r value was calculated across the r validation values. This process was repeated at 12 ridge regressions (λ values, 1×10^−3^:1×10^8^) with the lowest λ value, where any increase gave no further improvement to the average r value, was taken (47). Choosing the correct lambda value here again mitigated the potential overfitting of the TRF model. This average r value, at the optimal λ, was used for all further analysis.

### 2.6 mTRF auditory stimuli preprocessing

The envelope of the auditory signal was extracted by taking the absolute value of the analytic signal generated by the Hilbert transform (Matlab). As the envelope of the lower frequencies is linearly relatable to the EEG signal (16, 17, 53) the envelope of the stimuli was filtered between 0.5 Hz and 15 Hz (lowpass; 6^th^ order Butterworth filter. Highpass; 9^th^ order Butterworth filter). The resultant envelopes were normalised using *nt_normcol* (Noisetools). Finally, the stimulus envelopes were down sampled to 100 Hz to match the EEG signal.

### 2.7 mTRF Statistics

Random permutation statistics were created for each participant to measure the average stimulus reconstruction (r) that could be obtained by chance. The random permutation procedure was conducted per participant for each frequency band producing a paired chance stimulus reconstruction (r) per participant. To obtain a random permutation of the data, whilst maintaining phase integrity, each of the stimulus envelopes were first reversed and a random circular shift was applied. Next, the mTRF cross-validation was ran in the same way as the real data (see above for details), to give a stimulus reconstruction (r) value. This procedure was iterated 100 times to create a null distribution and the average of these 100 iterations were used as that participant’s random stimulus reconstruction (r) value.

### 2.8 mTRF Linear mixed effects model (LMEM)

A linear mixed effects model (LMEM), as described in the main text, was conducted in SPSS (IBM SPSS statistics 24), submitting data type (real r values, random permutation r values), frequency band (delta, theta or alpha) and age (4-, 7- or 11-months) along with interactions between data type by frequency band, data type by age and age by frequency band, to the LMEM. Due to the sensitivity of LMEM to outliers in the sample, outlier analysis was conducted to exclude data more than 1.5 interquartile ranges above the upper quartile or below the lower quartile separately per band and per age group. Following exclusion data remained as follows: delta (4-months N=53, 7-months N=53 and 11-months N=51), theta (4-months N=53, 7-months N=52 and 11-months N=55) and alpha (4-months N=53, 7-months N=52 and 11-months N=52). These were the data sets included in the LMEM (Table 2 and Figure 3). The Satterthwaite approximation was applied to approximate the degrees of freedom, due to the missing cases. Visual inspection of residual plots did not reveal any obvious deviations from homoscedasticity or normality.

**Table 1.**
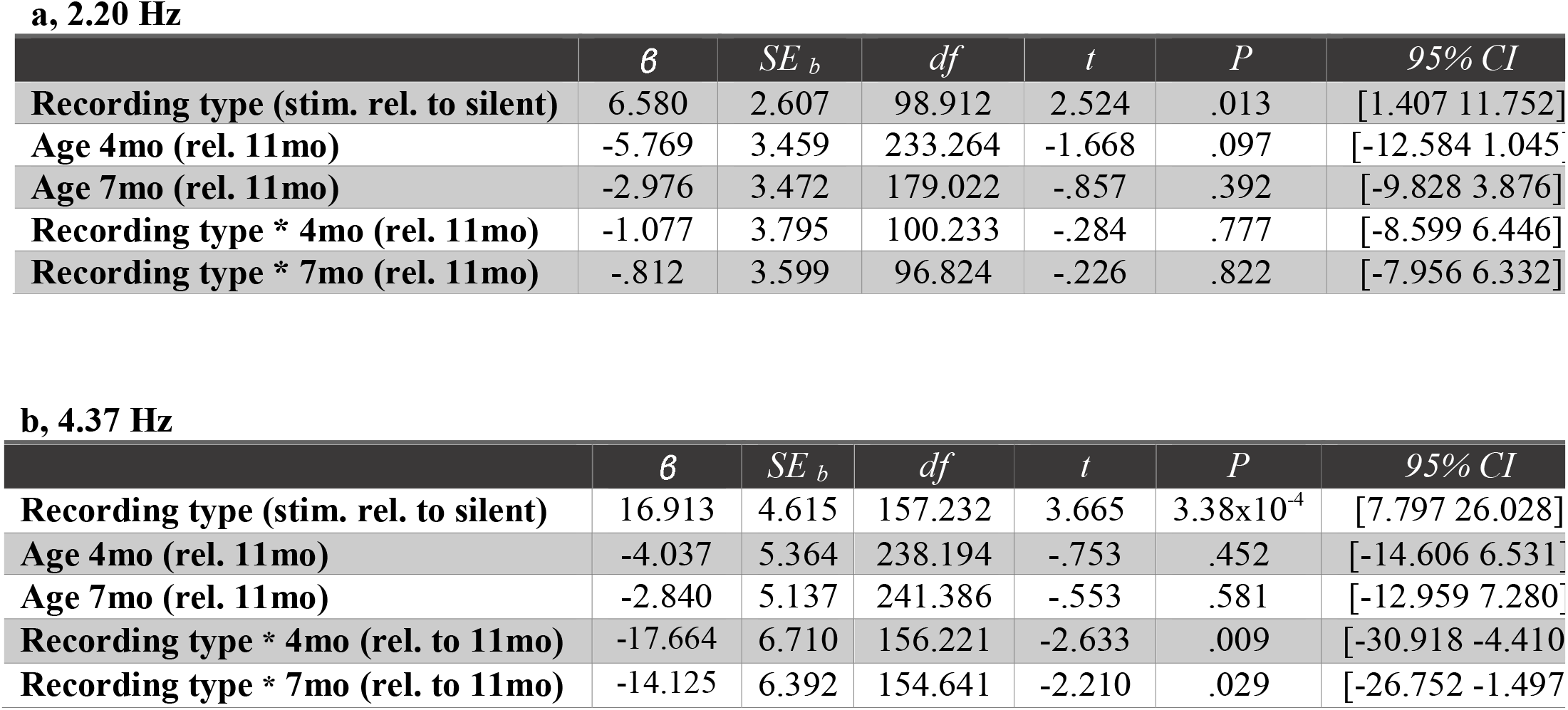
Parameter information detailing estimates of fixed effects on periodogram PSD peaks at a, 2.20 Hz and at b, 4.37 Hz, taken from the LMEM described in the text. The base cases for the LMEM were set as 11-months for age and silent state for recording type.

**a, 2.20 Hz**
| | $\beta$ | $SE_b$ | $df$ | $t$ | $P$ | 95% CI |
| --- | --- | --- | --- | --- | --- | --- |
| **Recording type (stim. rel. to silent)** | 6.580 | 2.607 | 98.912 | 2.524 | .013 | [1.407 11.752] |
| **Age 4mo (rel. 11mo)** | -5.769 | 3.459 | 233.264 | -1.668 | .097 | [-12.584 1.045] |
| **Age 7mo (rel. 11mo)** | -2.976 | 3.472 | 179.022 | -.857 | .392 | [-9.828 3.876] |
| **Recording type \* 4mo (rel. 11mo)** | -1.077 | 3.795 | 100.233 | -.284 | .777 | [-8.599 6.446] |
| **Recording type \* 7mo (rel. 11mo)** | -.812 | 3.599 | 96.824 | -.226 | .822 | [-7.956 6.332] |

**b, 4.37 Hz**
| | $\beta$ | $SE_b$ | $df$ | $t$ | $P$ | 95% CI |
| --- | --- | --- | --- | --- | --- | --- |
| **Recording type (stim. rel. to silent)** | 16.913 | 4.615 | 157.232 | 3.665 | $3.38 \times 10^{-4}$ | [7.797 26.028] |
| **Age 4mo (rel. 11mo)** | -4.037 | 5.364 | 238.194 | -.753 | .452 | [-14.606 6.531] |
| **Age 7mo (rel. 11mo)** | -2.840 | 5.137 | 241.386 | -.553 | .581 | [-12.959 7.280] |
| **Recording type \* 4mo (rel. to 11mo)** | -17.664 | 6.710 | 156.221 | -2.633 | .009 | [-30.918 -4.410] |
| **Recording type \* 7mo (rel. to 11mo)** | -14.125 | 6.392 | 154.641 | -2.210 | .029 | [-26.752 -1.497] |

**Table 2.** Parameter estimates from the LMEM. Individual subject person correlations (r values) derived from 3 frequency bands (delta 0.5-4 Hz, theta 4-8 Hz and alpha 8-12 Hz), 3 ages (4-, 7- or 11-months) and 2 data types (random or real). Random permutation r values, alpha and 11-months data were set as the respective base cases data type, frequency band and age in the model.

| | $\beta$ | $SE_b$ | $df$ | $t$ | $P$ | 95% CI |
| --- | --- | --- | --- | --- | --- | --- |
| **Data type (real rel. to rand for alpha base case)** | $-9.48 \times 10^{-4}$ | .001 | 901.802 | -.816 | .415 | [-.003 .001] |
| **Delta (rel. to alpha)** | .010 | .001 | 904.662 | 7.531 | $1.22 \times 10^{-13}$ | [.007 .012] |
| **Theta (rel. to alpha)** | .005 | .001 | 902.980 | 4.005 | $6.70 \times 10^{-5}$ | [.003 .008] |
| **Age (4mo rel. 11mo)** | -.002 | .001 | 907.156 | -1.365 | .173 | [-.004 $7.63 \times 10^{-4}$] |
| **Age (7mo rel. 11mo)** | $-2.84 \times 10^{-4}$ | .001 | 886.784 | -.221 | .825 | [-.003 .002] |
| **Data type \* delta (rel. to alpha)** | .012 | .001 | 901.802 | 9.549 | $1.19 \times 10^{-20}$ | [.010 .015] |
| **Data type \* theta (rel. to alpha)** | .004 | .001 | 901.802 | 3.039 | .002 | [.001 .006] |
| **Data type \* 4mo (rel. to 11mo)** | .003 | .001 | 901.802 | 2.471 | .014 | [.001 .006] |
| **Data type \* 7mo (rel. to 11mo)** | $6.36 \times 10^{-4}$ | .001 | 901.802 | .501 | .617 | [-.002 .003] |
| **4mo \* Delta (rel. to 11mo)** | .005 | .002 | 903.729 | 3.391 | .001 | [.002 .008] |
| **4mo \* Theta (rel. to 11mo)** | .001 | .002 | 902.587 | .803 | .422 | [-.002 .004] |
| **7mo \* Delta (rel. to 11mo)** | .001 | .002 | 903.407 | .901 | .368 | [-.001 .004] |
| **7mo \* Theta (rel. to 11mo)** | $8.70 \times 10^{-5}$ | .002 | 902.608 | .056 | .955 | [-.003 .003] |

The LMEM can be described by the following equation.

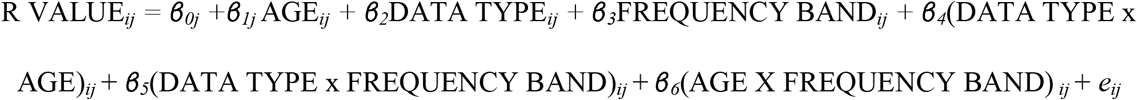

### 2.9 Phase amplitude coupling (PAC)

All remaining epochs after preprocessing were concatenated back into one continual EEG signal. Due to the increased susceptibility of PAC analysis to noise, manual rejection was conducted to further remove noisy periods of data from the signal.

To asses PAC, a modified version of the WinPACT plugin (EEGLab) (52) was used to acquire normalised modulation index (nMI) values (54), a measure adapted from Canolty and colleagues’ modulation index (MI) (10). The normalised version of the MI calculation (nMI) was selected as differences in low frequency power have been shown to adversely affect the PAC calculation (54–56). The MI method combines the amplitude envelope time series *A*_1_(*t* + τ) of a high-frequency with the phase time series φ_2_(*t*) of a specified low-frequency, creating a composite complex-valued signal *z*(*t*, τ). The resulting value is a widely validated metric of the coupling strength and preferred phase between two frequencies. For each infant’s EEG data, low-frequency phase (LFP) and high-frequency amplitude (HFA) were extracted with a zero-phase FIR filter (Hamming window), separately for all 60 electrodes. LFP centre frequencies were filtered from 2 to 8 Hz, in 1 Hz steps with a 2 Hz bandwidth, and HFA centre frequencies were filtered from 17.5 Hz to 42.5 Hz, in 5 Hz steps with a 5 Hz bandwidth. A sliding 5 second analysis window was implemented, with 2.5 second overlaps, with a mean vector length calculated per window. Next, 200 surrogate statistical iterations were created for each PAC calculation window. The statistically normalised MI estimate was obtained for each analysis window by subtracting the mean and dividing by the standard deviation obtained from a Gaussian fit of surrogate MI estimates (nMI = (Canolty’s MI – surrogate MI Mean) / surrogate MI Std). This statistical procedure was first suggested by Canolty et al. (2006) and implemented in the winPACT plugin based on code adapted from Özkurt and Schnitzler (54). Each iteration of the surrogate data was created by shuffling the high frequency amplitude time series via circular rotation. A MI estimate was obtained for each of the 200 surrogate data iterations, from which a 95% confidence interval was calculated using normcdf.m. This step accounted for the mean and standard deviation of the surrogate data set, thus creating an appropriate threshold for the frequency band analysed (see winPACT_precompute.m subscript in the winPACT toolbox for complete code, implemented in our analysis script ‘PAC_AA_final.m’ (54)). Finally, generalized family-wise error rate correction was implemented to correct for the multiple PAC calculation windows. The remaining statistically significant MI windows were averaged per channel for each of the PAC pairs (i.e. each LFP and HFA step) separately for each participant. The channel exhibiting the strongest MI, within predefined phase and amplitude band groupings (delta/beta, delta/gamma, theta/beta, theta/gamma), was taken forward for the LMEM and for the group level grand average plots.

### 2.10 PAC Linear mixed effects model

A LMEM was conducted using SPSS (IBM SPSS statistics 24), to investigate whether the statistically significant normalised MI estimates (nMI) (10, 54) were significantly different when either delta or theta was the low-frequency carrier phase, when either beta or gamma was the high-frequency amplitude and if this relationship was affected by the age of the infants. Fixed effects of low-frequency phase (LFP; delta or gamma), high-frequency amplitude (HFA; beta or gamma), and age (4-, 7- or 11-months) along with interactions between HFA by age, LFA by age and LFP by HFA were submitted to the LMEM. Due to the sensitivity of LMEMs to outliers in the sample, outlier analysis was conducted to exclude data more than 1.5 interquartile ranges above the upper quartile or below the lower quartile. Data sets remaining were 4-months N=54, 7-months N=52 and 11-months N=55 and these were included in the LMEM Table 3 and Figure 4. The Satterthwaite approximation was applied to approximate the degrees of freedom due to the missing cases.

**Table 3.** Parameter estimates from the LMEM. The normalised modulation index (nMI) measure of PAC was calculated between multiple low frequency phases (LFP) and High frequency amplitude (HFA) calculation pairs (LFP steps 2:8 Hz and HFA steps from 15:45 Hz). The maximum nMI value was taken from four pre-defined PAC band groupings (delta/beta, delta/gamma, theta/beta and theta/gamma; beta 2-4 Hz, theta 4-8 Hz, beta 15-30 Hz and gamma 30-45 Hz) and was submitted to the model. The base cases for the LMEM were set as 7-months for age, delta (*δ*) for LFP and beta (β) for HFA.

| | $\beta$ | $SE_b$ | $df$ | $t$ | $P$ | 95% CI |
| --- | --- | --- | --- | --- | --- | --- |
| **Low-frequency phase (rel. to $\delta$)** | .018 | .110 | 511.020 | .170 | .865 | [-.194 .231] |
| **High-frequency amplitude (rel. to $\beta$)** | .279 | .108 | 511.020 | 2.585 | .010 | [.067 .492] |
| **Age (4mo rel. to 7mo)** | .159 | .115 | 622.569 | 1.378 | .169 | [-.068 .385] |
| **Age (11mo rel. to 7mo)** | .148 | .115 | 586.146 | 1.288 | .198 | [-.078 .373] |
| **LFP \* 4mo (rel. to 7mo)** | -.013 | .132 | 511.020 | -.097 | .923 | [-.271 .246] |
| **LFP \* 11mo (rel. to 7mo)** | .080 | .131 | 511.020 | .612 | .541 | [-.177 .338] |
| **HFA \* 4mo (rel. to 7mo)** | -.209 | .132 | 511.020 | -1.585 | .114 | [-.468 .050] |
| **HFA \* 11mo (rel. to 7mo)** | -.250 | .131 | 511.020 | -1.907 | .057 | [-.508 .008] |
| **HFA \* LFP** | -.005 | .107 | 511.020 | -.043 | .965 | [-.215 .205] |

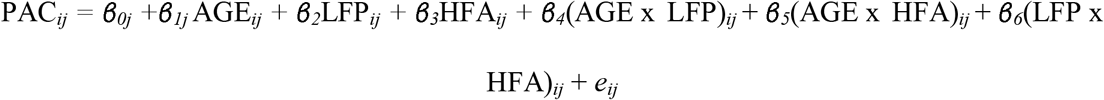

The LMEM can be described by the equation above, in which LFP denotes the band of the low-frequency phase (delta; 2-4 Hz or theta; 4-8 Hz), HFA is the band of the high-frequency amplitude (beta; 15-30 Hz or gamma; 30-45 Hz) and age is the age grouping of the infant (∼4-months, ∼7-months or ∼11-months). To best highlight the relative contribution of each level of the fixed effects in the beta weights, delta, beta and 7-months data were defined as the respective base cases.

### 2.11 Spectral analysis (periodogram PSD estimate)

The same concatenated data sets created for the PAC analysis were also used for the periodogram Power Spectral Density (PSD) estimates, in which all epochs after preprocessing were concatenated back into one continual signal with manual rejection used to remove noisy periods of data from the signal. A one-sided PSD estimate was conducted separately for each electrode channel using the periodogram function (Matlab). The length of the participants data was zero padded to ensure the size of the rectangular window used was equal in length to the number of discrete Fourier transform (DFT) points, ensuring the correct FFT across participants. This resulted in 52,834 equal spaced frequency bins from 0 to 50 Hz.

The periodogram can be defined by the following formula. In which the EEG signal, *x*_n_, is sampled at 100 Hz, with Δ*t* as the sampling interval.

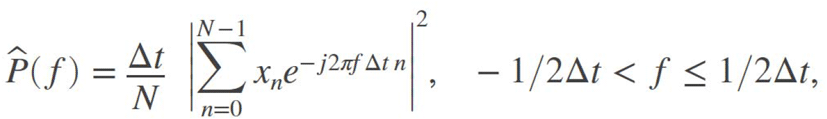

To achieve the one-sided periodogram output reported in Figure 1, values at all frequencies (except 0 and the Nyquist, 1/2Δt), were multiplied by two to conserve the total power.

**Figure 1.**
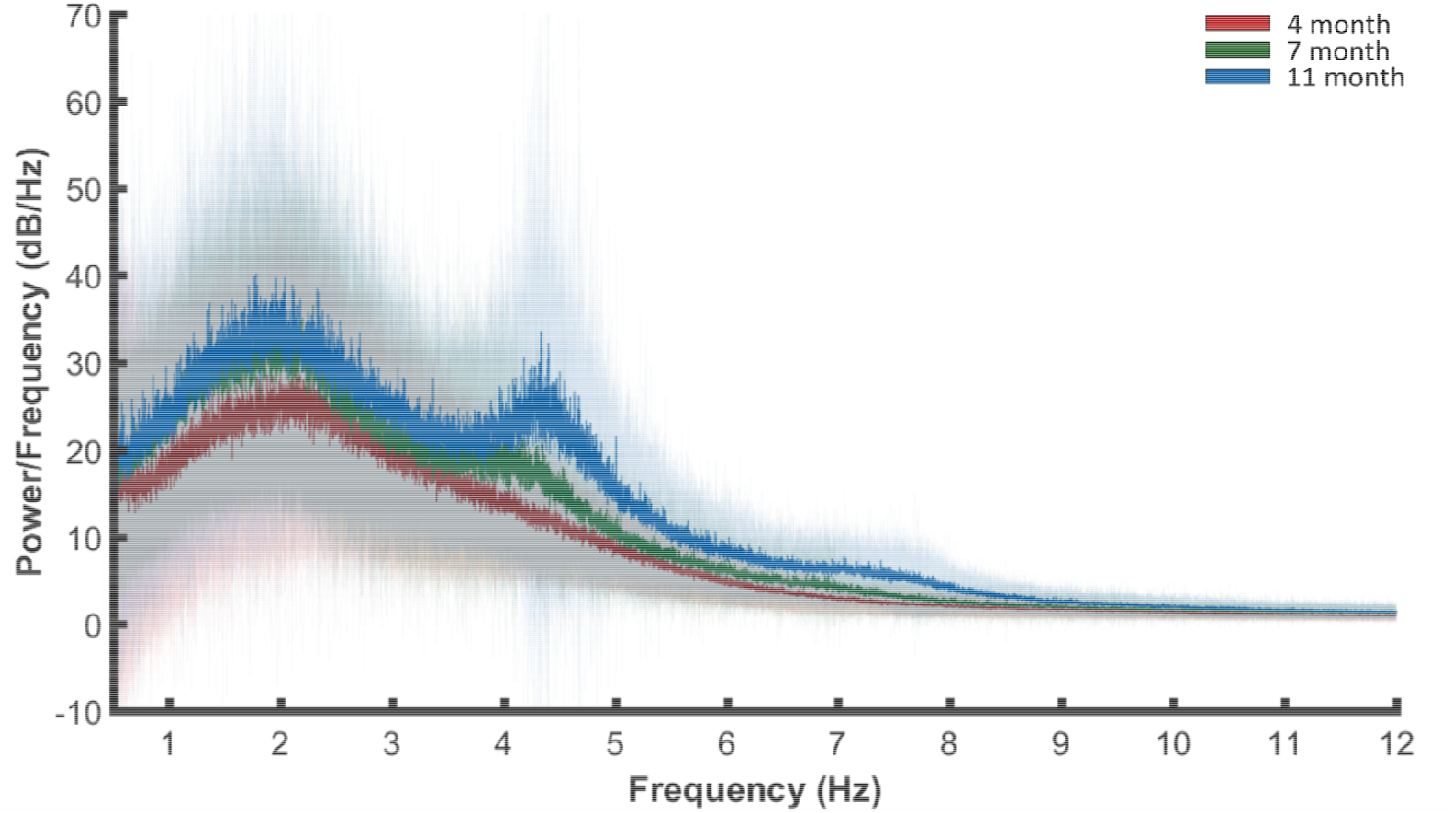
Spectral decomposition of the EEG signal (0.5-12 Hz plotted from 0.5-50 Hz calculated) in response to nursery rhyme stimulation. A periodogram was used to obtain a power spectral density (PSD) estimate separately for 4- (red), 7- (green) and 11- (blue) months data. Bold lines indicate the mean values and pale shading plots the standard deviation of the data. Outlier analysis was also conducted to remove extreme data points that would compromise the LMEM (detailed rejection criteria given in methods). The data plotted included, N=44 4-month, N=47 7-month and N=50 11-month old infants.

### 2.12 PSD Linear mixed effects model

To investigate whether the PSD was significantly higher in response to the nursery rhymes compared to the silent state, and whether this relationship was affected by the age of the participants, a linear mixed effects model (LMEM) was conducted separately for the 2.20 Hz and 4.37 Hz frequency peaks (IBM SPSS statistics 24). To allow for individual subject variation, peak values per participant were taken as the maximum value of a participants’ 60 channel average data, within a 1 Hz window centered around the peak of interest (2.20 Hz or 4.37 Hz). Fixed effects of recording type (nursery rhymes, silent state), age (4-, 7- or 11-months) and a recording type by age interaction were included in the model. Silent state data and 11-months data were set as the respective base cases for each variable in the model. Due to the sensitivity of LMEMs to outliers in the sample, outlier analysis was conducted to exclude data more than 1.5 interquartile ranges above the upper quartile or below the lower quartile. The final number of data sets included in the LMEM were; stimulus period (4-months N=44, 7-months N=47, 11-months N=50) and silent state (4-months N=35, 7-months N=46, 11-months N=38). These data are also used to generate Figure 1. The Satterthwaite approximation was applied to approximate the degrees of freedom due to the missing cases.

The model can be described by the following equation:

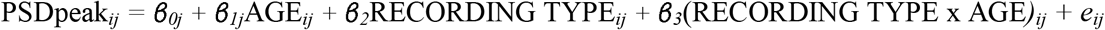

## 3 Results

### 3.1 Both delta and theta EEG frequency bands respond to nursery rhymes

Prior to investigating cortical tracking and PAC, the distribution of low-frequency neural signals within our data was established using spectral decomposition of the signal, achieved using the periodogram power spectral density (PSD) estimate. After preprocessing, PSD was obtained for each of the remaining 60 electrodes in response to audio-visually presented nursery rhymes and during a five minute period of silence. A grand average across channels and ages revealed two frequency peaks centered around 2.20 Hz and 4.37 Hz for the nursery rhyme period (Supplementary Figure S1). These peak responses corresponded to the frequency peaks found in the acoustic envelope of the nursery rhymes (Supplementary Figures S2, S3 and S4). Separating the data by age revealed ∼2.20 Hz PSD peaks were consistently present at 4-, 7- and 11-months (Figure 1), whereas, the mean values of the ∼4.37 Hz peaks displayed an age-related increase in PSD.

To investigate whether the PSD was significantly higher in response to the nursery rhymes compared to the silent state, and whether this relationship was affected by the age of the participants, a linear mixed effects model (LMEM) was conducted separately for the 2.20 Hz and 4.37 Hz frequency peaks (see methods). The average PSD across all 60 electrodes was used, with the peak value (within the 1 Hz window centered on the peak of interest) submitted to the model for each participant. Fixed effects of recording type (nursery rhyme stimulus period, silent state), age (4-, 7- or 11-months) and a recording type by age interaction were included in the model. Silent state data and 11-months data were set as the respective base cases for each effect in the model. Individual differences across the 3 recording sessions (at 4-, 7- and 11-months) were accounted for by including a random intercept (AR1) per participant.

For the 2.20 Hz peak, tests of fixed effects revealed that recording type had a significant effect (*F*(1, 98.560) = 15.513, *P* = 1.53×10^−4^) on PSD, as the ∼2.20 Hz PSD increased significantly during the stimulus period (*M* = 29.782, *SE* ± 1.670) vs the silent state (*M*=23.832, *SE* ±1.751). Neither age (*F*(2, 137.925) = 2.780, *P* = 0.066) nor age by recording type (*F*(2, 98.427) = 0.045, *P* = 0.956) showed an effect on PSD. Accordingly, the stimulus effect present at 11-months is also present at 4- and 7-months (see Table 1a).

For the 4.37 Hz peak, recording type again showed a significant effect (*F*(1, 154.963) = 5.560, *P* = 0.020) on PSD, increasing during the stimulus period (*M* = 21.428, *SE* ± 2.180) vs silent state (*M*=15.111, *SE* ± 2.341). Furthermore, both age (*F*(2, 201.747) = 6.406, *P* = 0.002) and age by recording type (*F*(2, 154.883) = 4.005, *P* = 0.020) exerted a significant effect on PSD. Exploration of the estimates of fixed effects showed that the significant interaction between recording type by age was driven by a greater effect at 11-months (Table 1b). Bonferroni-corrected pairwise comparisons showed that PSD increased significantly between 4- and 11-months (*M* increase = 12.869, *SE* ± 3.793, *P* = 0.002) and 7- and 11-months (*M increase* = 9.902, *SE* ± 3.783, *P* = 0.029). Hence there was a developmental increase in 4.37 Hz PSD for both the nursery rhymes and the silent state. This may indicate age-related maturation of theta activity. Importantly, we also observed a stimulus-driven interaction with age for the 4.37 Hz PSD peaks, with the largest increase observed at 11-months (silent state PSD peak, *M* = 17.404, *SE* ± 3.909 vs nursery rhymes PSD peak, *M* = 34.316, *SE* ± 3.437). This shows that the interaction was driven by a greater PSD in response to the nursery rhymes at 11-months.

The data indicate that the nursery rhymes produced a ∼2 Hz PSD response as early as 4-months of age whereas the nursery rhymes produced a ∼4 Hz response that developed between 4- and 11-months.

### 3.2 Delta and theta EEG frequency bands track nursery rhyme envelopes

Power spectral density (PSD) analysis confirmed that the nursery rhyme stimuli showed increased power in the delta (∼2.20 Hz) and theta (∼4.37 Hz) bands. To investigate the presence and strength of cortical tracking, backward mTRFs (Figure 2a) (47) were trained with either delta (0.5-4 Hz), theta (4-8 Hz) or alpha (8-12 Hz) EEG signals. The decoding model was fit separately to the Hilbert envelope of each of the 83 nursery rhyme trials, for each participant, using a leave-one-out cross-validation procedure (see methods). Pearson’s correlation (r) was used to test the quality of the reconstruction (Figure 2b). This provided an objective metric of envelope tracking at the individual level. To test the correlation (r) values against chance, random permutation statistics were created for each participant (N = 100 permutations).

**Figure 2.**
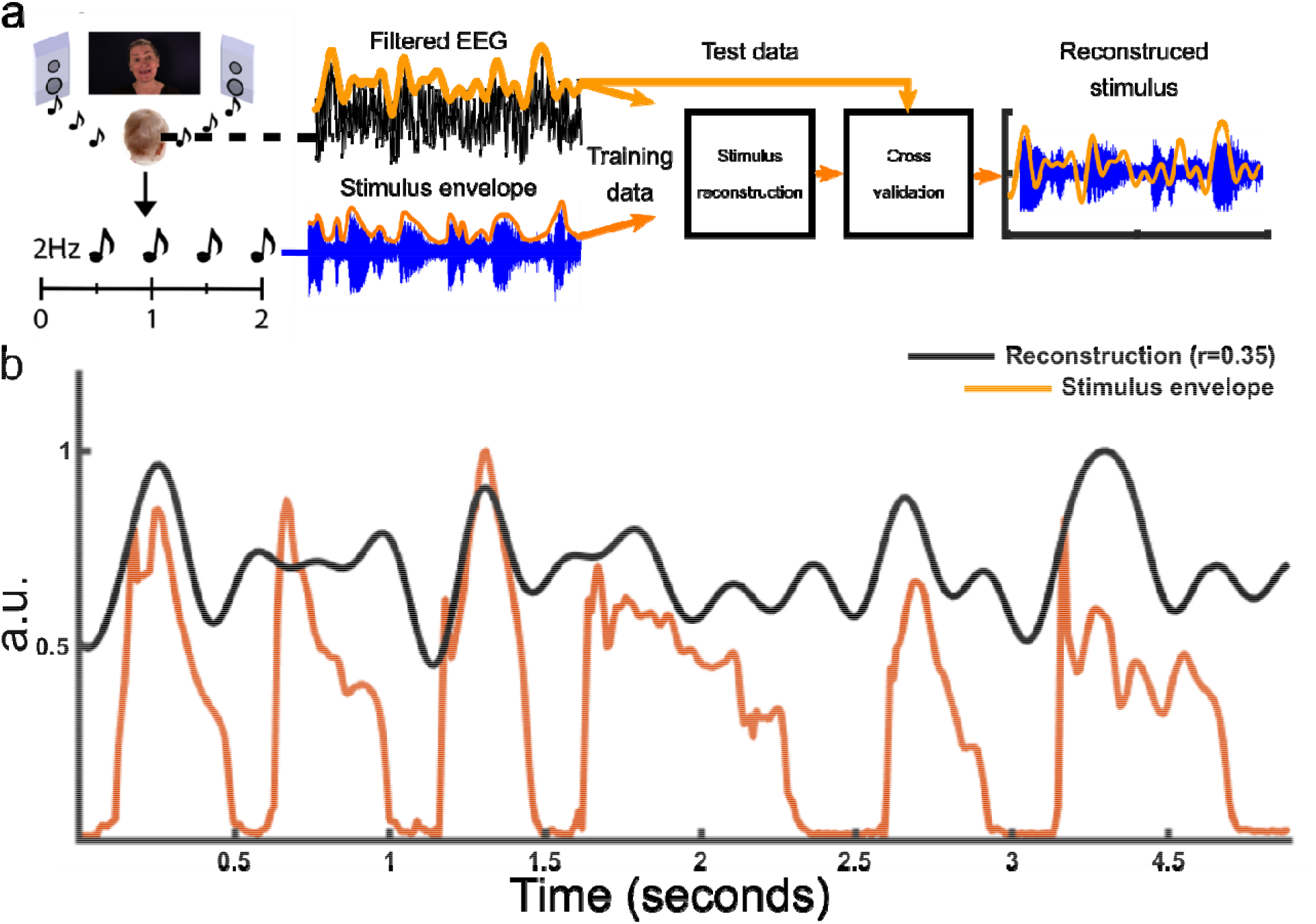
mTRF stimulus reconstruction. a) Schematic of the experimental procedure along with a summary of the mTRF analysis pipeline. The EEG signal and the stimulus envelope (absolute value of the Hilbert envelope) were submitted to the mTRF stimulus reconstruction. For the cross validation procedure, 83 nursery rhyme trials were rotated M-1 times each serving once as the “test set” with the remainder of the trials being the “training set”. The process was repeated at 12 ridge regressions (λ values, 1×10^−3^:1×10^8^) with the average model convolved with the test data to reconstruct the stimulus envelope at the optimal λ. b) Example of one of the 83 mTRF stimulus reconstructions for one participant along with the original acoustic stimulus envelope. The black line depicts the reconstruction (in arbitrary units) and the orange line illustrating the absolute value of the Hilbert envelope of the stimulus (in arbitrary units).

The reconstruction analyses showed that cortical delta and theta neural signals significantly tracked the envelopes of the nursery rhyme stimuli, but alpha signals did not (Figure 3).

**Figure 3.**
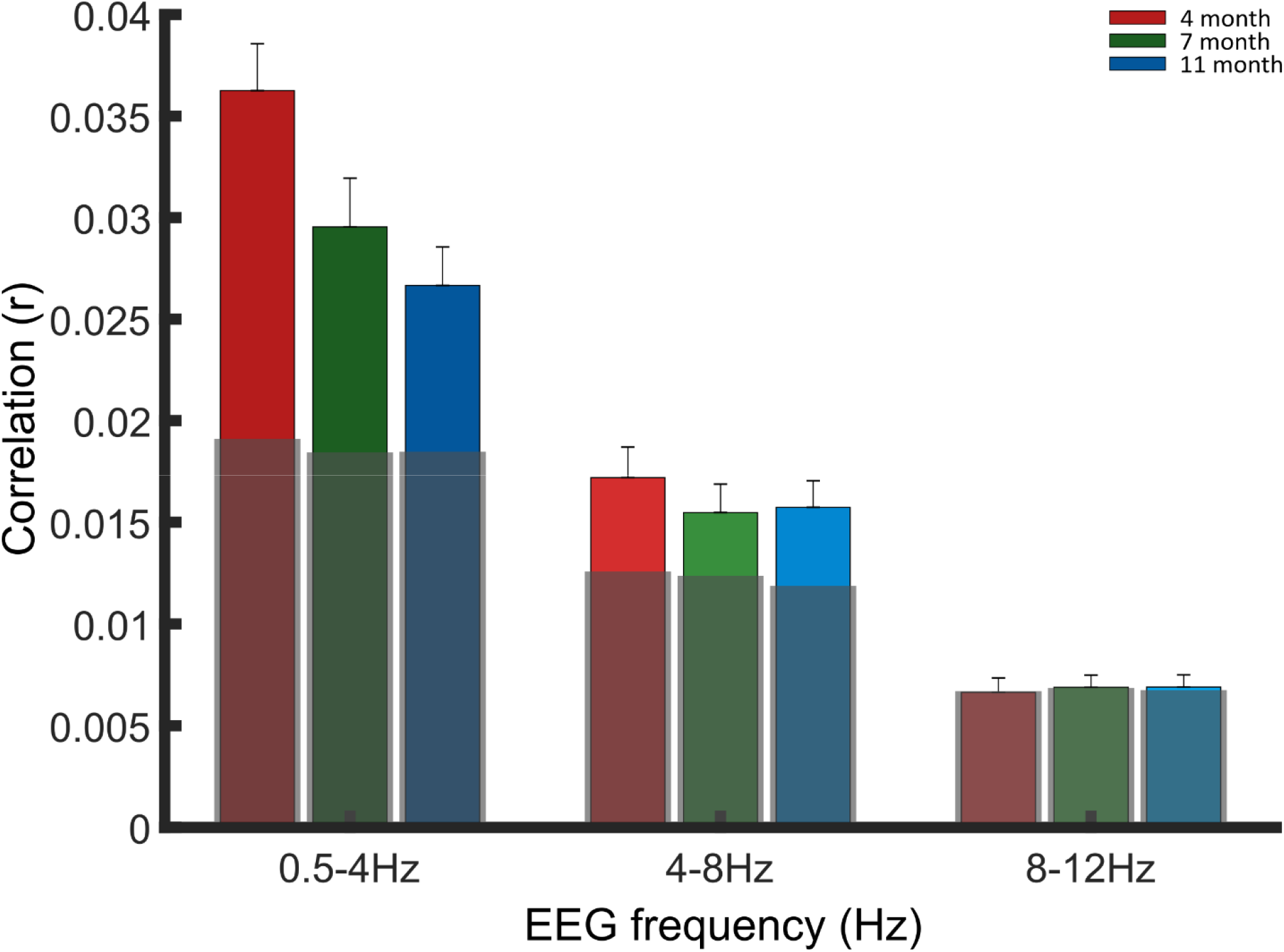
Grand averaged stimulus reconstruction values (Pearson’s r) per frequency band. Each bar represents the grand average r value across the participants at the 3 age points; 4-months (red), 7-months (green) and 11-months (blue). Age responses are grouped along the X axis by EEG frequency band, delta (0.5-4 Hz) theta (4-8 Hz) and alpha (8-12 Hz). Light grey boxes signify the average random permutation value across subjects. Outlier analysis was conducted to remove extreme values which would compromise the LMEM, full rejection numbers detailed in methods.

A LMEM was conducted to investigate potential developmental changes in cortical tracking. Main effects of data type (real r values, random permutation r values), frequency band (delta, theta or alpha) and age (4-, 7- or 11-months) were investigated along with interactions between data type by frequency band, data type by age and age by frequency band. Random permutation r values, alpha frequency band and 11-month data were set as the respective base cases for each variable in the model. A random intercept (AR1) per participant was included to account for individual differences across the 3 recording sessions (4-, 7- and 11-months).

Tests of fixed effects showed a significant effect of data type (*F*(1, 901.802) = 118.656, *P* = 4.806 x10^−26^), as the real data values were significantly higher than the random permutation values (Figure 3). A significant effect of frequency band (*F*(2,903.216) = 401.939, *P* = 1.399 x10^−125^) highlighted that cortical tracking (r values) differed by frequency band, with the estimates of fixed effects confirming greater tracking in both delta and theta bands compared to the alpha base case (Table 2). Furthermore, estimates of fixed effects for the alpha base case (Table 2) confirmed that alpha band itself did not produce significant cortical tracking above the randomly permuted values. A significant interaction between data type and frequency band (*F*(2,901.802) = 47.657, *P* = 2.114 x10^−20^) indicated that cortical tracking in the delta and theta bands was driven by their values being significantly above the randomly permuted r values (as compared to the alpha base case; Table 2). A significant effect of age (*F*(2,755.514) = 5.079, *P* = 0.006) showed that the strength of the cortical tracking values changed with age. There were significant interactions between age by data type (*F*(2,901.802) = 3.413, *P* = 0.033) and age by frequency band (*F*(4,902.774) = 3.372, *P* = 0.009). Inspecting the estimates of fixed effects (Table 2), the data shows that the significant difference between random permutation and real r values is strongest in the 4 month data and the frequency band differences between the age groups is predominantly driven by greater cortical tracking in the delta band at 4 months (as compared to 11 months).

To highlight the relative contribution of delta and theta to the LMEM, the model’s base case within frequency band was changed from alpha to theta. Estimates of fixed effects revealed that delta produced significantly larger cortical tracking (r values) than theta (Supplementary Table S1). This increase was again driven by the real rather than the random permutation data.

The mTRF stimulus reconstruction data show that delta and theta EEG responses track the acoustic envelope of sung and chanted speech from the age of 4-months, supporting our hypothesis that delta-band cortical tracking is developmentally important. Furthermore, 4-month infants showed significantly greater tracking than older infants, and this was strongly influenced by the strength of the delta-band response. This could either reflect bottom-up infant use of acoustic landmarks at a delta-band rate in the sung and chanted speech, facilitating attentional mechanisms such as grouping, or could reflect the fact the speech was sung and therefore rhythmic (57), or both. Theta-band neural signals also tracked the speech envelope, but at a significantly lower level. In contrast to the developmental maturation observed in theta power (Figure 1), there was no evidence for a developmental change in theta envelope tracking during the first year of life. Significant theta tracking was observed as early as 4-months, the age at which infants are thought to recognize their first word (58) (their own name). In accordance with our sensory hypothesis (51), as infants become more efficient speech comprehenders, top-down processes may be expected to increase the degree of theta tracking beyond that observed here. Increased attention to speech content with increasing comprehension may also introduce alpha tracking, perhaps during the second year of life.

### 3.3 Phase Amplitude Coupling (PAC)

PAC was measured using the normalised Modulation Index (nMI) method (54), in which frequency band specific normalisation was applied to Canolty’s Modulation Index (MI) calculation (10). This measure combines the amplitude envelope of a selected high-frequency band with the phase time series of a low-frequency band, creating a complex-valued signal referred to as the MI. The MI estimate of PAC was selected as it has been shown to be the most tolerant to signal noise (59), a common issue in infant EEG. It was important to apply the normalised version of the MI calculation (nMI) as differences in low frequency power have been shown to adversely affect the PAC calculation (54–56). Taking the nMI value allowed us to directly compare the role of delta vs theta phase with a more robust estimate than MI itself (54).

For each participant low-frequency phases from 2 to 8 Hz (1 Hz steps) and high-frequency amplitudes from 15 to 45 Hz (5 Hz steps) were extracted from the EEG signal from each of the 60 electrode channels. For each of these PAC pairing steps, multiple nMI values were calculated per infant via a 5-second sliding window. The significant windows were identified if they exceeded the 95% confidence interval calculated from a surrogate data set made up of 200 statistical iterations of the same analysis window (see methods for full details). The nMI estimate was obtained by utilizing the surrogate data (10, 54) by subtracting the mean and dividing by the standard deviation (obtained from a Gaussian fit of all the surrogate MI estimates) from each MI value. All analysis windows that produced significant nMI estimates, within each PAC pairing, were next averaged per participant. To allow for individual subject variation, the PAC pairing with the maximum nMI value from within predefined phase and amplitude band groupings (delta/beta, delta/gamma, theta/beta, theta/gamma) was taken, before a grand average (Figure 4) was calculated for each phase and amplitude band grouping (see methods).

**Figure 4.**
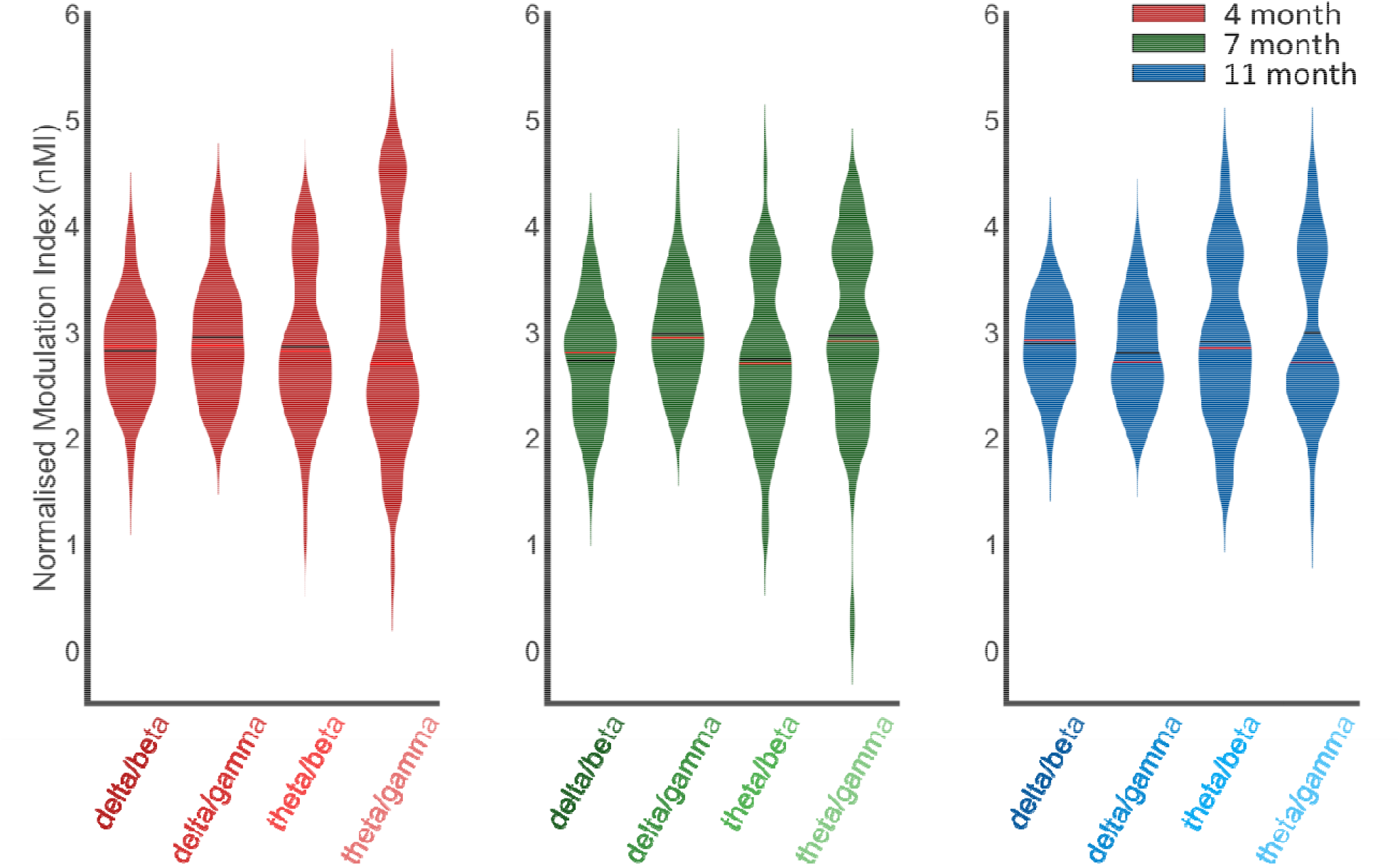
Violin plots of phase amplitude coupling (normalised modulation index, nMI) at 4-months (red), 7-months (green) and 11-months (blue). The nMIs, from all significant analysis windows, were averaged for each individual infant. The PAC pairing with the maximum nMI per infant, from within the pre-defined frequency bands of interest (delta/beta, delta/gamma, theta/beta and theta/gamma; beta 2-4 Hz, theta 4-8 Hz, beta 15-30 Hz and gamma 30-45 Hz), were included in the grand average violin plot. Horizontal black and red lines represent the group mean and median respectively. Outlier analysis was conducted to remove extreme data points that would compromise the LMEM, therefore data plotted consisted of the following number of data points (4-months, N=54; 7-months N=52; 11-months N=55).

To investigate whether the nMI values were significantly different when either delta or theta was the low-frequency carrier phase or when either beta or gamma were the high-frequency amplitudes, and if this relationship was affected by the age of the infants, a LMEM was conducted. Fixed effect factors of low-frequency phase (delta or theta), high-frequency amplitude (beta or gamma) and age (4-, 7- or 11-months) were inputted to the LMEM. The maximum nMI value from the four PAC band pairings (delta/beta, delta/gamma, theta/beta and theta/gamma), from pre-defined frequency band groupings (delta 2-4 Hz, theta 4-8 Hz, beta 15-30 Hz and gamma 30-45 Hz) were extracted per participant and submitted to the model. Interactions of low-frequency phase by age, high-frequency amplitude by age and low-frequency phase by high-frequency amplitude were also included in the model, with a random intercept (AR1) per participant included to account for individual differences across the 3 ages of recording (see methods). Delta, beta and 7-months data were defined as the respective base cases.

Tests of fixed effects showed a significant effect only for high-frequency amplitude (*F*(1, 511.020) = 5.404, *P* = 0.020). Bonferroni-corrected pairwise comparisons showed MI values were larger when gamma (*M* = 2.952, *SE* ± .040) rather than beta (*M* = 2.827, *SE* ± 0.040) was the high-frequency amplitude. Tests of fixed effects were not significant for low-frequency phase (*F*(1, 511.020) = .522, *P* = 0.470), age (*F*(2, 316.195) = 0.468, *P* = 0.626), age by low-frequency phase (*F*(2, 511.020) = 0.301, *P* = 0.740), age by high-frequency amplitude (*F*(2, 511.020) = 2.070, *P* = 0.127), nor low-frequency phase by high-frequency amplitude (*F*(1, 511.020) = 0.002, *P* = 0.965).

In summary, delta and theta both acted as significant carrier phases, coupling with higher-frequency amplitudes in the infant brain. Whilst both beta and gamma amplitudes significantly coupled with delta and theta low frequency phases, the nMI values were significantly greater when gamma rather than beta was the HFA (*Mean diff*. = 0.125, *SE* ± 0.040).

## 4 Discussion

Here we show, for the first time, that infant delta and theta cortical signals track the rhythmic fluctuations of the speech envelope of sung IDS. We demonstrate that delta tracking of the speech envelope is significantly stronger than theta tracking during the first year of life, and that delta tracking is particularly strong at the earliest measurement, 4 months. The stimulus-produced delta signal power (PSD) is constant from 4-months, whilst theta power develops continuously between 4- and 11-months. Also for the first time, we demonstrate PAC in response to speech, at all ages studied. Contrary to our initial hypothesis, we show that delta acts as an equally strong carrier phase of higher-frequency neural signals as theta. The data thus suggest that both delta and theta play important roles in the temporal organisation of higher frequency amplitudes in the infant brain. As also found in the adult literature, high frequency gamma amplitudes seem to couple with delta and theta to a greater extent than beta amplitudes. However, PAC to the same sung speech in adults has not yet been analysed, to test whether similar PAC patterns are observed (Attaheri 2021; in preparation). By contrast to adult studies, alpha cortical tracking was not observed for infants in the current paradigm. This is not surprising. As our participants are pre-verbal, top-down mechanisms related to language encoding and attentional prediction of speech should not yet be present. (28, 60). The likelihood that our infants were not yet comprehending the nursery rhymes may explain the absence of alpha tracking, as alpha tracking may reflect both attention to and comprehension of speech (61).

Significant cortical tracking was already evident in our sample at 4-months of age. Indeed, non-speech studies with newborn infants (using amplitude-modulated noise) have shown that delta cortical tracking of rhythm is present at birth (62). Accordingly, language experience may not be required for efficient delta-band cortical tracking of speech rhythm (51). The strong and early delta-band tracking observed here may provide a foundation for the attentional and grouping mechanisms that support perceptual organization of the speech signal, as indicated by adult studies (24). Here, rather than reflecting the use of discourse-level information, it is likely that the acoustic landmarks provided by stressed syllable placement in the varied nursery rhymes underpin the dominant delta-band tracking observed. However, delta-band tracking may also reflect the speech input being sung and thus highly rhythmic, although note that a range of different beat rates and rhythmic patterns (trochaic, iambic, dactyl) were utilized for the current nursery rhyme corpus (35). To disentangle these possibilities, similar studies with adults are required, which we are currently engaged in (Attaheri 2021; in preparation). Animal studies have also demonstrated cortical tracking of rhythmically-structured acoustic input in both delta and theta bands (9, 63), possibly suggesting that these are general auditory perceptual abilities conserved across mammalian species. Nevertheless, animals do not learn spoken language, while both IDS and nursery rhymes are highly rhythmic and optimally-structured to support infant language learning (46). IDS has increased delta-band modulation energy and significantly stronger delta-theta phase synchronization of AMs than ADS (64). Accordingly, it seems plausible that infants, who are exposed to maternal speech rhythms while in the womb, may use these general perceptual abilities to entrain to human speech. Following birth, they may rapidly augment low-frequency cortical tracking with the higher-frequency information perceived by the human ear, for example via PAC. Given the importance of canonical babbling in the first year of life, the delta-beta PAC reported here may indicate the importance of motor prediction of speech for the language-learning brain (65). Early infant babbling is known to reflect the rhythmic properties of the adult language, as behavioural studies show that the babble of Arabic-, French- and Cantonese-learning infants are distinguishably different (66).

It has also been demonstrated in adult studies that adding more acoustic features (such as spectral features) to the envelope information studied here improves the performance of TRF models. In a similar vein, it seems plausible that the infant brain may rapidly augment basic delta-band tracking with a rich repertoire of other linguistic elements as language comprehension grows. It is known from fMRI studies that by age 3 months the infant cortex is already structured into several regions of functional importance for speech processing, for example Broca’s area and anterior temporal regions (67). While the neural mechanisms underlying synchronization with the auditory envelope during speech listening are actively debated in the adult literature (1), there is gathering evidence that the alignment of cortical neural signals to specific stimulus parameters of speech plays a key role in language processing and comprehension (6, 17, 53). Accordingly, the cortical tracking approach employed here will enable us to see whether individual differences in cortical tracking predict individual differences in language outcomes. We are currently collecting these language data. If individual differences in cortical tracking do predict individual differences in language outcomes, it would support the view that the infant brain synchronises with attended speech in order to learn language. As our sample ages, we plan to investigate these possibilities.

## Author Contributions

*Adam Attaheri*: EEG Paradigm Development, EEG Preprocessing, Investigation - Data Curation, Formal Analysis - design, creation and implementation of analysis, Writing - Original Draft.

*Áine Ní Choisdealbha*: EEG Preprocessing - Investigation - Data Curation - Writing - Review & Editing

*Giovanni Di Liberto*: Formal analysis, Writing - Review & Editing

*Sinead Rocha:* Investigation - Data Curation, Writing - Review & Editing

*Perrine Brusini*: EEG Paradigm Development, Investigation - Data Curation

*Natasha Mead*: Investigation - Data Curation

*Helen Olawole-Scott*: Investigation - Data Curation

*Panagiotis Boutris*: Investigation - Data Curation

*Samuel Gibbon*: Investigation - Data Curation

*Isabel Williams*: Investigation - Data Curation

*Christina Grey*: Investigation - Data Curation

*Sheila Flanagan*: Investigation - Modulation analysis

*Usha Goswami*: Conceptualization - Methodology, Funding Acquisition, Supervision, Project Administration, Writing - Original Draft

## Data Sharing agreement

The analyses were conducted by using publicly available MATLAB toolboxes that can be downloaded at http://audition.ens.fr/adc/NoiseTools/, https://sccn.ucsd.edu/eeglab/download.php, https://github.com/mickcrosse/mTRF-Toolbox. The custom code used for the mTRF, PSD and PAC analysis can be downloaded from https://osf.io/q9s5k/. The final data included in the manuscript figures and statistics can also be downloaded from https://osf.io/q9s5k/.

This data is currently being analysed with other techniques as part of the ongoing longitudinal BabyRhythm project. Once this research has been completed, the raw EEG files will be uploaded to the same OSF server as above. In the meantime, the raw data can be made available upon request.

## Acknowledgements

We would like to thank all the families and infants who kindly donated their time to this project. We would also like to thank Henna Ahmed for her time spent as an RA on the BabyRhythm project. Final we would like to acknowledge Mike X Cohen’s statistical textbooks and online materials for guiding our statistical analysis.

