## Supplemental figures and tables for "Delta- and theta-band cortical tracking and phase-amplitude coupling to sung speech by infants"

**Supplement**

**Grand average PSD across 4-, 7- and 11-months in response to nursery rhymes**

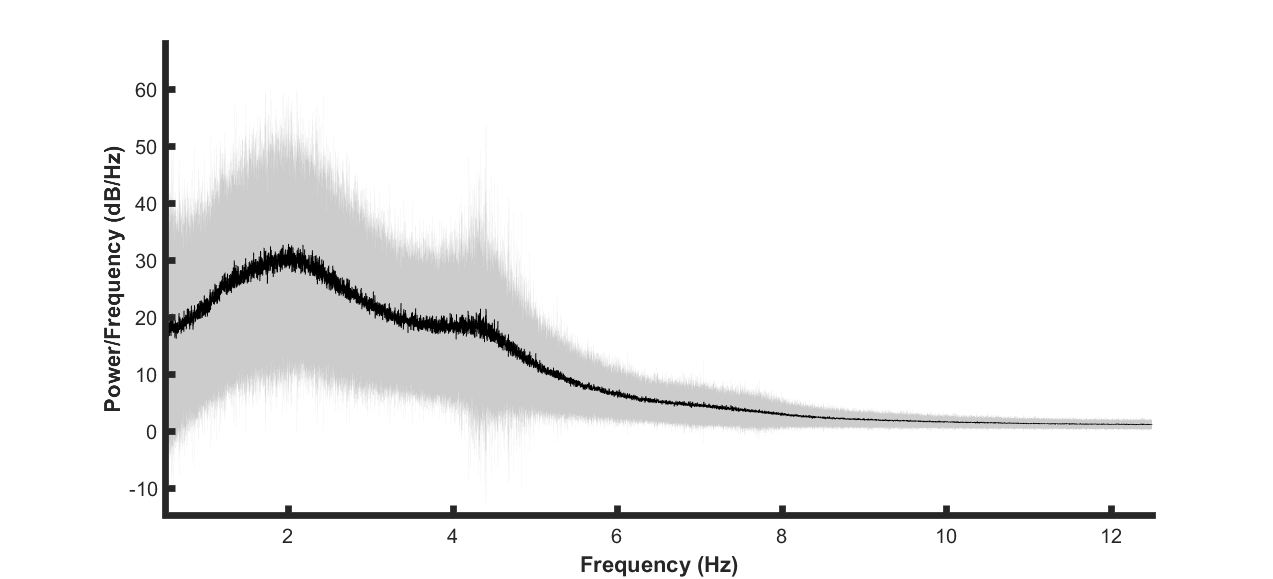

Figure S1, Spectral decomposition of the EEG signal (0.5-12 Hz). A periodogram was used to obtain a power spectral density (PSD) estimate separately for 4-, 7- and 11-months data, before being averaged together to create a grand average. The bold black line indicates the mean values and pale shading plots the standard deviation of the data. Outlier analysis was also conducted to remove extreme data points that would compromise the LMEM (detailed rejection criteria in main methods). The data plotted are for nursery rhyme stimulation only (4-months N=44, 7-months N=47, 11-months N=50).

**Modulation Spectrum of nursery rhyme sound files**

All analysis was computed using MATLAB 2016a. The modulation spectrum extraction was based on an approach described by Plomp, et al., (1) and developed by Leong, et al. (2). First, the sound files were down sampled to 14.7 kHz and then band-pass filtered using a series of adjacent FIR filters, into five bands: 100-300 Hz, 300-700 Hz, 700-1750 Hz, 1750-3900 Hz, and 3900- 7250 Hz. Next, the Hilbert envelope was extracted from each of the five sub-band signals. The five envelopes were down sampled to 1050 Hz then filtered through a modulation filter bank. This modulation filter bank comprised 24 channels logarithmically spaced between 0.9-40 Hz. In Figure S2, the RMS difference power was averaged across 18 nursery rhymes and determined for each spectral band. In Figure S3, for one nursery rhyme, the RMS power in each spectral band was divided by the overall RMS power, revealing the relative amount of energy in each band. In Figure S4, the “all band averages” from each of the 18 individual nursery rhymes are depicted together, as denoted by the thin multicolored lines. The grand average of these 18 individual “all band averages” and STD are denoted by the black line and grey shading respectively. Modulation filter bank corner frequencies were taken as [0.93; 1.09; 1.27; 1.49; 1.74; 2.03; 2.38; 2.78; 3.25; 3.80; 4.45; 5.20; 6.08; 7.11; 8.32; 9.72; 11.38; 13.30; 15.56; 18.20; 21.28; 24.89; 29.11; 34.04; 39.81]. Clear peaks in modulation power can be observed at ~2.18 and ~4.4Hz.

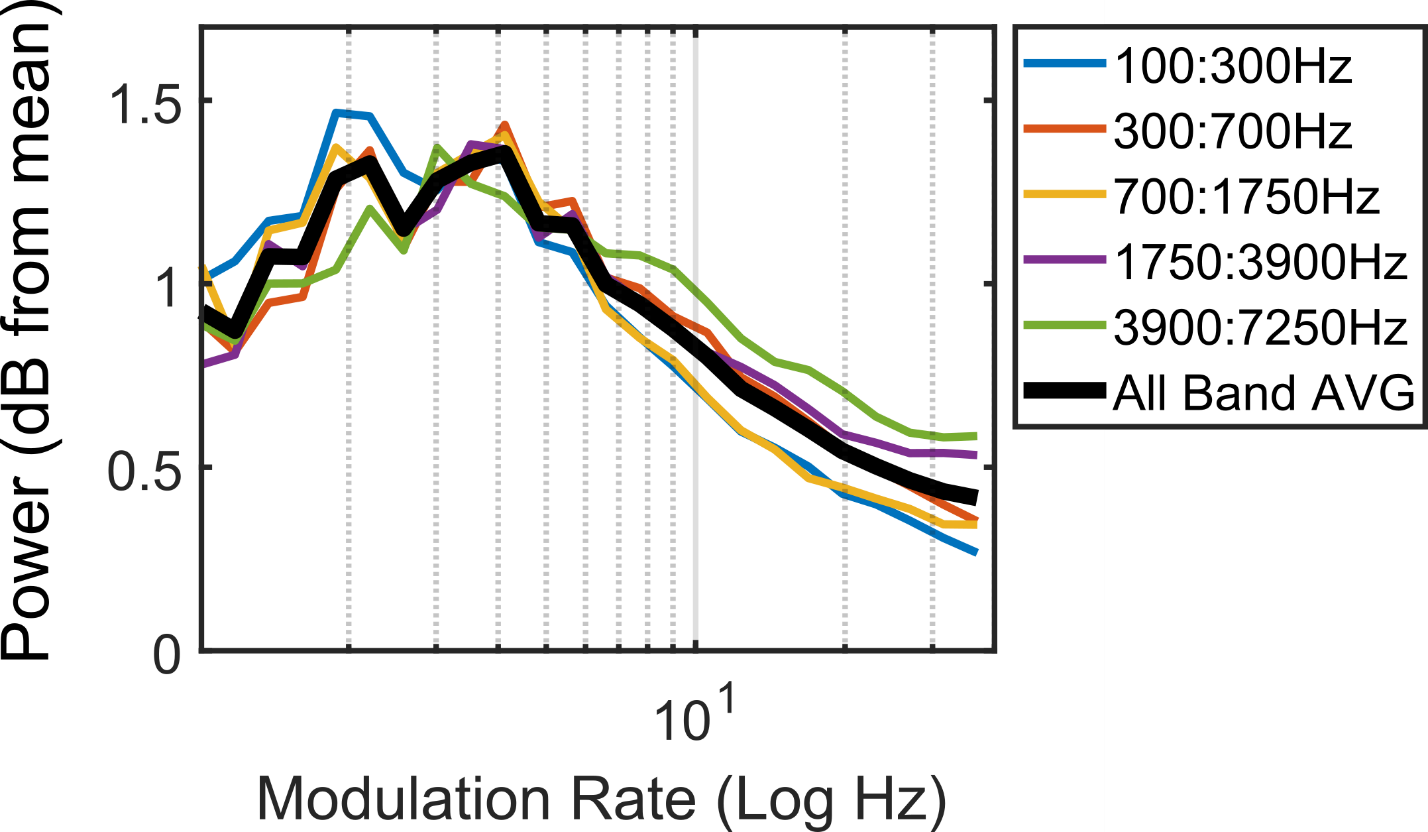

Figure S2, Average modulation spectrum of all the Nursery rhyme stimuli.

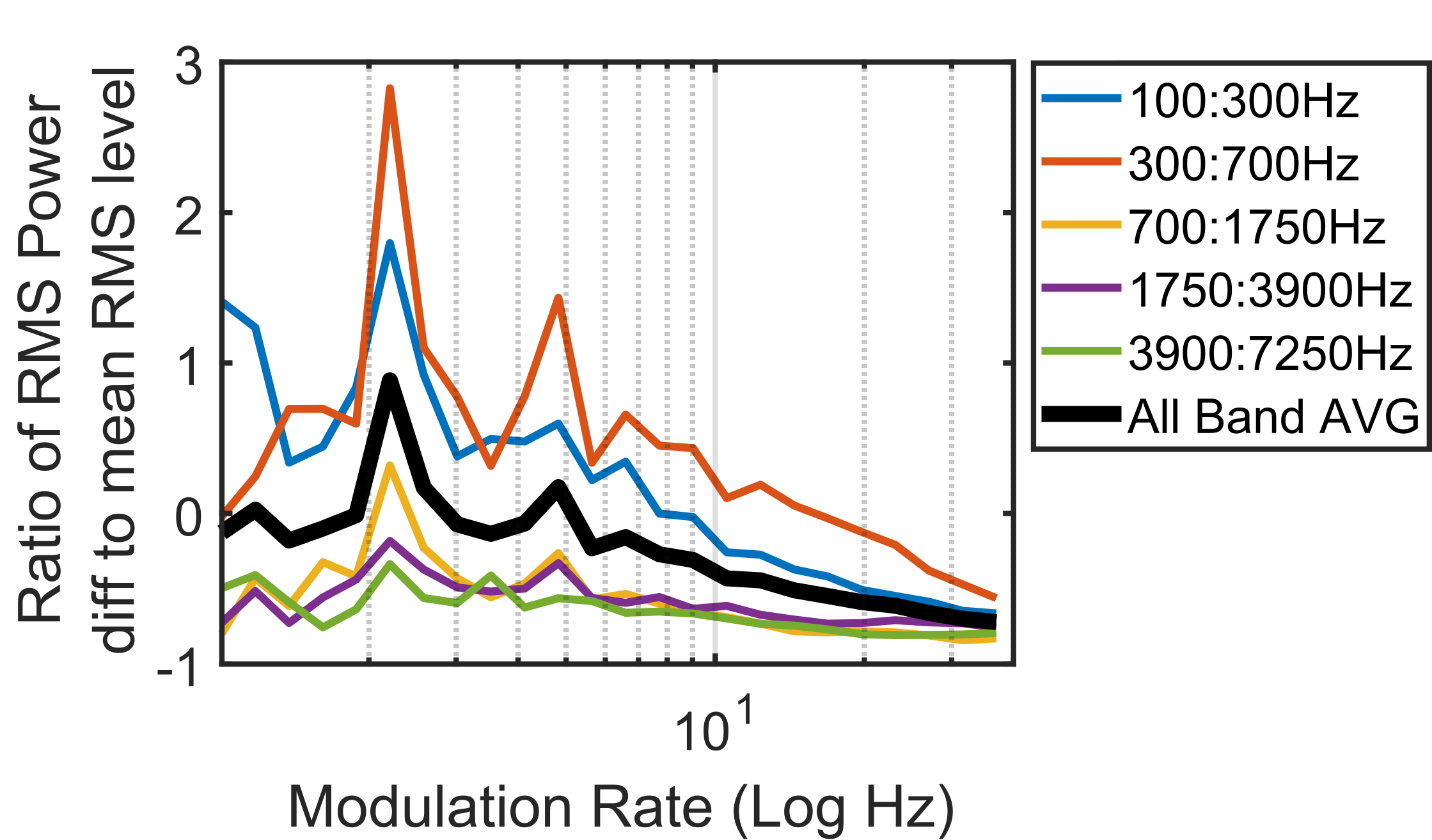

Figure S3, Example Modulation spectrum of one the Nursery rhyme stimuli (‘Simple Simon’). RMS power in each spectral band was derived by the overall RMS power.

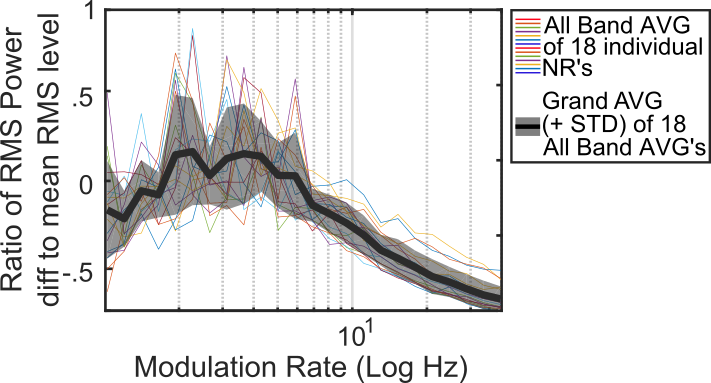

Figure S4, Grand average of the 18 individual nursery rhymes “All Band AVG” Modulation spectrums, with the “All Band AVG” depicted for each individual rhyme by the separate multicolored lines. The grand average of the 18 individual “All Band AVG”, and their standard deviations (STD), are denoted by the black line and grey shading respectively.

**Clean_ASR EEGLab function**

Artifact Subspace Reconstruction (ASR) (3) is an automated artifact rejection function. It uses principle component analysis (PCA) to ensure no period of the data signal has abnormally strong power. First the algorithm finds the cleanest portion of the data to use as the calibration data, and the first statistics are computed. Next, a 0.5 second sliding window PCA (with 50% window) is performed across all the channels to identify bad PCs. The algorithm next finds the subspaces in which the activity that is more than 5 standard deviations away from the calibration data. The high variance subspaces (i.e. group of channels associated with the ‘bad’ PC’s) are then reconstructed using a mixing matrix that was calculated on the clean data.

**LMEM examining mTRF r values with theta base case for frequency band**

A Linear Mixed Effects Model (LMEM) was ran to highlight the relative contribution of delta and theta frequency bands, on the Pearsons correlation (r) values. This LMEM was constructed with the same factors as the LMEM described in the main text (main effects of data type [real r values, random permutation r values], frequency band [delta, theta or alpha] and age [4-, 7- or 11-months], along with interactions between data type by frequency band, data type by age and age by frequency band. However, to highlight the relative contribution of delta and theta, the base case for frequency band was changed to theta. The base cases for data type and age where kept as real r values and 11-months data respectively. A random intercept (AR1) per participant was included to account for individual differences across the 3 recording sessions (4-, 7- and 11-months). As the model remained the same, and only the base case was changed, the tests of fixed effects showed identical tests of fixed effects as to those found in the main text. Parameter estimates from the LMEM are given in Table S1.

| **LMEM estimates of fixed effects for Pearson correlation (r) values** | | | | | | |
| --- | --- | --- | --- | --- | --- | --- |
|  | *β* | *SE _b_* | *df* | *t* | *P* | *95% CI* |
| **Data type (real rel. to rand for theta base case)** | .003 | .001 | 901.802 | 2.531 | .012 | [6.52 x10^-10^ .005] |
| **Delta (rel. to theta)** | .005 | .001 | 903.367 | 3.630 | .005 | [.002 .007] |
| **Alpha (rel to theta)** | -.005 | .001 | 902.980 | -4.005 | .000 | [-.008 -.003] |
| **Age (4mo rel. to 11mo)** | -5.03x10^-4^ | .001 | 905.613 | -.398 | .691 | [-.003 .002] |
| **Age (7mo rel. to 11mo)** | -1.96x10^-4^ | .001 | 886.323 | -.155 | .877 | [-.003 .002] |
| **Data type * delta (rel. to theta)** | .008 | .001 | 901.802 | 6.554 | 9.41x10^-11^ | [.006 .011] |
| **Data type * alpha (rel. to theta)** | -.004 | .001 | 901.802 | -3.039 | .002 | [-.006 -.001] |
| **Data type * 4mo (rel. to 11mo)** | .003 | .001 | 901.802 | 2.471 | .014 | [6.44x10^-4^ .006] |
| **Data type * 7mo (rel. to 11mo)** | 6.36x10^-4^ | .001 | 901.802 | .501 | .617 | [-.002 .003] |
| **4mo * Delta (rel. to 11mo)** | .004 | .001 | 902.847 | 2.614 | .009 | [.001 .007] |
| **4mo * Alpha (rel. to 11mo)** | -.001 | .001 | 902.587 | -.803 | .422 | [-.004 .002] |
| **7mo * Delta (rel. to 11mo)** | .001 | .001 | 903.112 | .851 | .395 | [-.002 .004] |
| **7mo * Alpha (rel. to 11mo)** | -8.70x10^-5^ | .001 | 902.608 | -.056 | .955 | [-.003 .003] |

Table S1. Parameter estimates from the LMEM. Individual subject person correlations (r values) derived from 3 frequency bands (delta 0.5-4Hz, theta 4-8Hz and alpha 8-12Hz), 3 ages (4-, 7- or 11-months) and 2 data types (random or real). Random permutation r values, theta and 11-months data were set as the respective base cases data type, frequency band and age in the model.

**Random effects from the Linear Mixed Effects Models (LMEM)**

**PSD model**

At 2.18 Hz, the random intercept in the model demonstrated that the effect of age on PSD showed significant variance across participants (AR1 diagonal estimate = 89.364, *X^2^* (1) = 4.177, *P* = 2.9 x10^-5^ ) and this variance across participants significantly covaried between each repeated measure (AR1 rho estimate = -0.330, *X^2^* (1) = -2.246, *P* = .025 ).

At 4.40 Hz, the random intercept in the model demonstrated that the effect of age on PSD showed significant variance across participants (AR1 diagonal estimate = 75.491, *X^2^* (1) = 2.756, *P* = 0.006), however this variance across participants did not significantly covary between each repeated measure (AR1 rho estimate = -0.315, *X^2^* (1) = -1.876, *P* = .061 ).

**mTRF model**

The random intercept in the model demonstrated that the variance of age across participants did not show significant covariance between each repeated measure (AR1 rho estimate = .347, *X^2^* (1) = 0.732, *P* = .464).

**PAC model**

The random intercept in the mTRF model demonstrated that the effect of age on PAC (MI) did not show significant variance across participants (AR1 diagonal estimate = .0068, *X^2^* (1) = .885, *P* = .376) nor did the variance across participants significantly covary between each repeated measure (AR1 rho estimate = 0.097, *X^2^* (1) = .665, *P* = .885 ).

**Data storage**

Study data were collected and managed using REDCap electronic data capture tools hosted at Cambridge university (4). REDCap (Research Electronic Data Capture) is a secure, web-based software platform designed to support data capture for research studies, providing 1) an intuitive interface for validated data capture; 2) audit trails for tracking data manipulation and export procedures; 3) automated export procedures for seamless data downloads to common statistical packages; and 4) procedures for data integration and interoperability with external sources.
